# DAF-16/FOXO maintains genome integrity following genotoxic stress

**DOI:** 10.1101/2025.10.27.684284

**Authors:** Umanshi Rautela, Oviya Devendran, Gautam Chandra Sarkar, KR Ranjisha, Rashi Mittal, Anita Goyala, Arnab Mukhopadhyay

## Abstract

Preserving genomic integrity is crucial for the accurate transmission of genetic information across generations, as well as for preventing precocious ageing. The DNA Damage Response (DDR) safeguards the genome from genotoxic stress through a coordinated system of sensors, relay proteins, and repair mechanisms. Since DNA repair is an energy-intensive activity, the process is tightly regulated and coordinated with various metabolic pathways. The nutrient-sensing insulin/IGF signalling (IIS) pathway has been extensively studied for its role in ageing and lifespan regulation in *C. elegans* through its downstream FOXO transcription factor DAF-16. However, there is limited understanding of its involvement in maintaining genomic integrity through the regulation of the DDR. In this study, we demonstrate the role of DAF-16/FOXO in preserving genome integrity by activating the expression of DDR repair genes in *C. elegans*. Activated DAF-16/FOXO directly binds to the promoter of DDR genes under conditions of low IIS, ensuring that their expression is maintained at a higher level, which is crucial for prompt DNA damage repair. Interestingly, we find that DAF-16 functions both cell autonomously as well as non-autonomously to support DNA integrity. We also determine that the DAF-16(d/f) isoform, but not the DAF-16(a) isoform, is essential for maintaining germline genome integrity. Furthermore, DAF-16 activation enhances the DDR primarily through the canonical DDR components and, to a lesser extent, via apoptosis-mediated clearance of damaged cells. Overall, our study highlights a new role for DAF-16/FOXO in the DDR and the preservation of genome integrity.

## Introduction

Maintaining genome stability in the face of environmental and endogenous genotoxic stressors is essential for organismal survival, fertility, and longevity. The organism’s DNA damage response (DDR) comprises pathways that detect DNA lesions, halt cell cycle progression, and facilitate repair to prevent genome instability [1]. As DDR is an energetically demanding process, it is tightly linked to various metabolic pathways such as the pentose-phosphate pathway to provide nucleotide precursors [2] or the fatty acid oxidation-dependent DNA break detection via poly (ADP-ribose) polymerase 1 (PARP1) [3]. Additionally, NAD⁺ metabolism impacts the activity of PARPs and sirtuins, which are essential for chromatin relaxation and DNA strand break repair [4] [5]. Among key metabolic regulators, the nutrient-sensing insulin/IGF-1 signaling (IIS) pathway serves as a conserved modulator of stress resistance and lifespan [6]. Central to the IIS pathway is the FOXO transcription factor (TF), which remains cytoplasmic and inactive under normal/high insulin signaling, but translocates to the nucleus under low IIS to initiate the transcription of a wide array of protective genes [7–9]. While FOXO transcription factors are well known for their roles in metabolism, stress resilience, and lifespan extension [10], emerging studies over the past decade have identified them as important modulators of genome stability.

In mammalian systems, FOXO3a regulates cell-cycle checkpoint activation (p21/Kip-1 and p27/Cip-1) [11, 12], oxidative DNA damage repair [13] and also DNA damage-induced apoptosis [14–16]. FOXO3 is linked to enhanced DNA damage repair and has been shown to physically interact with the ataxia-telangiectasia mutated (ATM) protein kinase, a master regulator of DDR [17]. Additionally, FOXO is implicated in transcriptional regulation of Gadd45a (a growth arrest and DNA damage response gene)[18] and is known to influence varying modes of DDR pathways (BER: base excision repair; HR: Homologous recombination)[13, 19]. However, how the FOXO activation mechanistically contributes to the DNA repair efficiency has not been comprehensively studied.

The nematode *Caenorhabditis elegans* is a powerful model to study DDR due to its genetic tractability, conserved DDR pathways, and accessible germline, enabling real-time visualization and quantification of DNA damage responses. Importantly, the distinct utilization of homologous recombination (HR)-mediated DDR in the germline and non-homologous end joining (NHEJ) in somatic tissues provides a unique opportunity to investigate tissue-specific DDR mechanisms [20, 21]. To probe the role of insulin/IGF-1-FOXO signalling in DDR, we employed the temperature-sensitive kinase-domain mutant *daf-2(e1370),* which mimics low IIS and activates the FOXO ortholog DAF-16 [22–24].

In the current study, we show that DAF-16 directly regulates DDR gene expression and improves damage resolution. Using tissue-specific knockdown and isoform-rescue experiments, we found that DAF-16 acts both cell-autonomously and cell non-autonomously to maintain germline genomic integrity, and the DAF-16(d/f) isoform is specifically required for enhancing germline DDR. Finally, we demonstrate that the improved DDR in *daf-2* mutants is mediated by homologous recombination (HR)-mediated DNA repair rather than increased apoptotic clearance. Together, our findings reinforce the role of FOXO/DAF-16 as a guardian of genome integrity, mechanistically linking the metabolic status of the cell to DNA damage repair.

## Results

### DAF-16 directly regulates DNA damage response genes in *daf-2* mutants

The IIS pathway modulates longevity and stress resistance in *C. elegans*, largely through the FOXO transcription factor DAF-16. Under reduced IIS (as in *daf-2* mutants), DAF-16 translocates to the nucleus and activates target genes involved in stress resistance, metabolism, and lifespan extension [23–26]. While the role of DAF-16 in stress resistance and longevity [27] is well-established, its regulation of the DNA damage response (DDR) remains less explored.

To investigate whether DAF-16 influences DDR pathways, we analyzed transcriptomic data from late-L4 staged *daf-2* and *daf-16;daf-2* mutants (previously published transcriptomics data) [28]. Strikingly, DDR genes were significantly upregulated in *daf-2* mutants compared to *daf-16;daf-2* double mutants (**Fig. 1A**), suggesting that their induction depends on DAF-16. We validated these findings by qRT-PCR, confirming elevated expression of key DDR genes such as *rad-50, rad-51, brc-1, hus-1* in the *daf-2* mutants (**Fig. 1B**). Moreover, in comparison to the wild-type (WT) worms, we observed an increased expression in the basal levels of DDR gene expression in *daf-2* worms (**Fig. S1A**).

**Fig 1:**
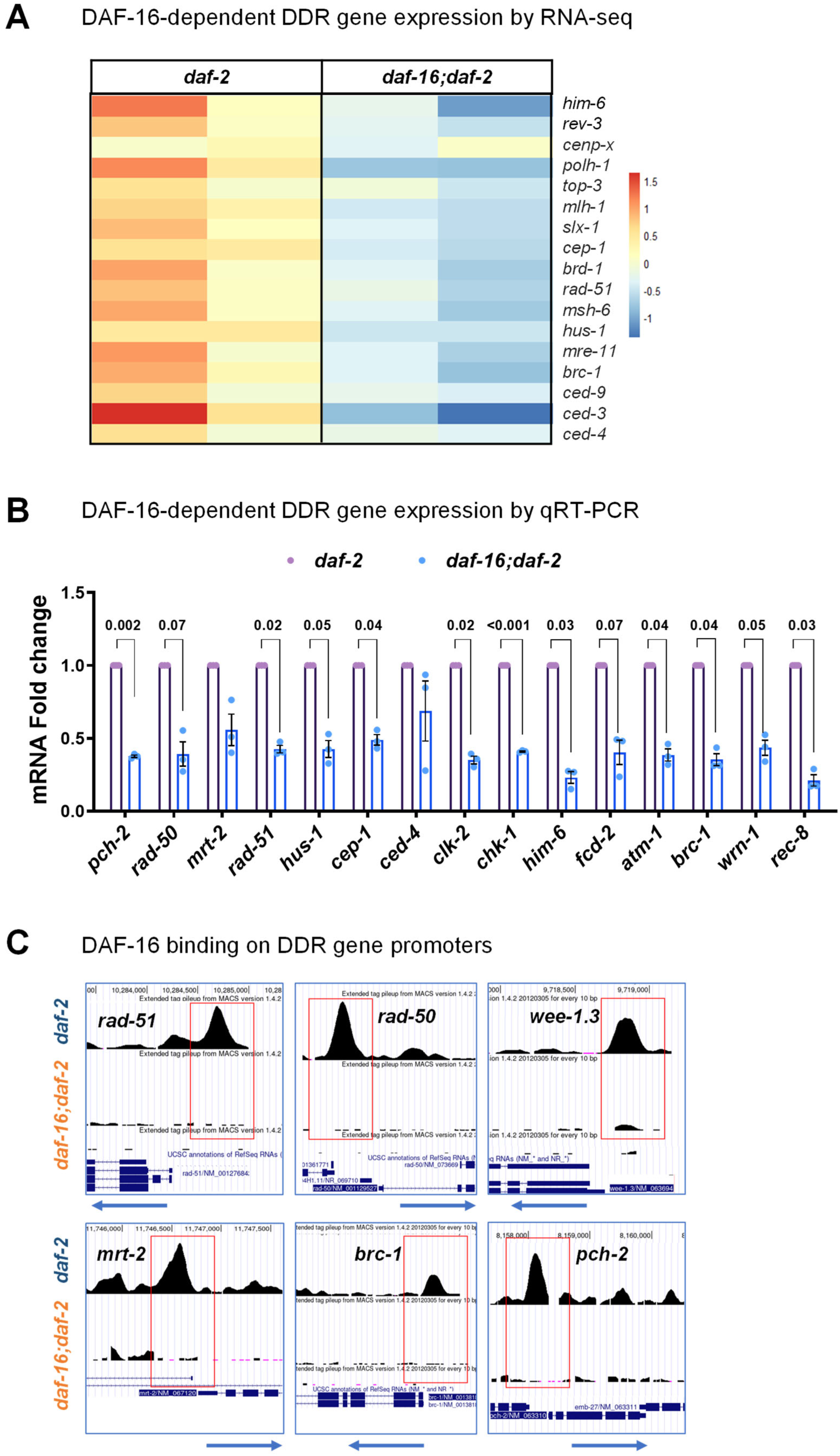
DAF-16/FOXO regulates DNA damage repair pathway gene expression. **(A)** Heat map representation of the levels of DNA damage repair (DDR) genes from transcriptomics data comparing *daf-2(1370)* and *daf-16(mgdf50);daf-2(e1370)* late-L4 staged worms grown on control RNAi. **(B)** Quantitative RT-PCR analysis showing mRNA fold change in DDR gene expression in *daf-2(1370)* and *daf-16(mgdf50);daf-2(e1370)* late-L4 staged worms grown on control RNAi. Expression levels were normalized to *actin*. The average of three biological replicates is shown. Unpaired *t*-test with Welch’s correction. **(C)** UCSC browser view of DAF-16/FOXO peaks on the DDR gene (*rad-51, rad-50*, *wee-1.3*, *mrt-2, brc-1* or *pch-2)* promoters as determined by ChIP-seq using an anti-DAF-16 antibody [29]. Red boxes indicate the binding on promoter regions of DDR genes. The upper panel shows *daf-2(e1370)*, where DAF-16 peaks are observed, while lower panel shows *daf-16(mgDf50);daf-2(e1370)* (that lacks specific DAF-16 peaks). Experiments were performed at 20°C. Source data are provided in Table 1.

Since DAF-16 is predominantly inactive in WT worms but nuclear-localized in *daf-2* [23], we hypothesized that DAF-16 may directly regulate the DDR genes. To test this, we re-analyzed published DAF-16 ChIP-seq data [29] and identified strong DAF-16 binding peaks at the promoters of multiple DDR genes in *daf-2* (**Fig. 1C**). These peaks were absent in *daf-16;daf-2* mutants, indicating that DAF-16’s binding is specific and likely drives the transcriptional activation.

Together, these results demonstrate that DAF-16, already known to govern stress and longevity pathways, also directly regulates DDR genes under reduced IIS, potentially linking DNA repair mechanisms to the extended lifespan of *daf-2*.

### DAF-16 Activation Enhances DNA Damage Repair in Germline and Somatic Tissues

In *C. elegans* hermaphrodites, oocytes pause at diakinesis, the final stage of meiotic prophase, prior to fertilization. At this stage, chromosomes are highly condensed, with homologous pairs no longer exhibiting the side-by-side pairing observed earlier in prophase. Instead, they remain physically linked through chiasmata formed by meiotic crossovers, enabling proper orientation on the meiotic spindle. In WT oocytes, six distinct DAPI-stained bivalents are typically visible, representing the six homologous chromosome pairs maintained by these chiasmata [30]

To investigate the role of DAF-16 in DDR, we exploited this well-defined chromosomal organization and performed a chromosome fragmentation assay. For this, worms were exposed to ionizing radiation (IR; dose of 0 Gy and 100 Gy) at the late-L4 stage and DAPI-stained 48 hours post-irradiation (HPI) to observe the extent of chromosome fragmentation in the oocytes [31]. Irradiation (100 Gy) of WT worms induced severe chromosomal fragmentation and fusion in diakinesis-stage oocytes, disrupting the normal bivalent structure. Strikingly, *daf-2* mutants, where DAF-16 is constitutively active, maintained intact chromosomal morphology, with clearly discernible bivalents (**Fig. S2A, B**). This protection was entirely DAF-16-dependent, as irradiated *daf-16;daf-2* double mutants exhibited extensive chromosomal defects (**Fig. 2A, B**). While at 100 Gy IR dose, both WT and *daf-16* worms displayed severe chromosomal abnormalities, at a reduced IR dose (70 Gy), *daf-16* mutants showed significantly more chromosomal aberrations (fragmentation and fusion) than WT (**Fig. S2C, D**). This demonstrates a role of DAF-16 in preserving meiotic chromosome integrity.

**Fig. 2:**
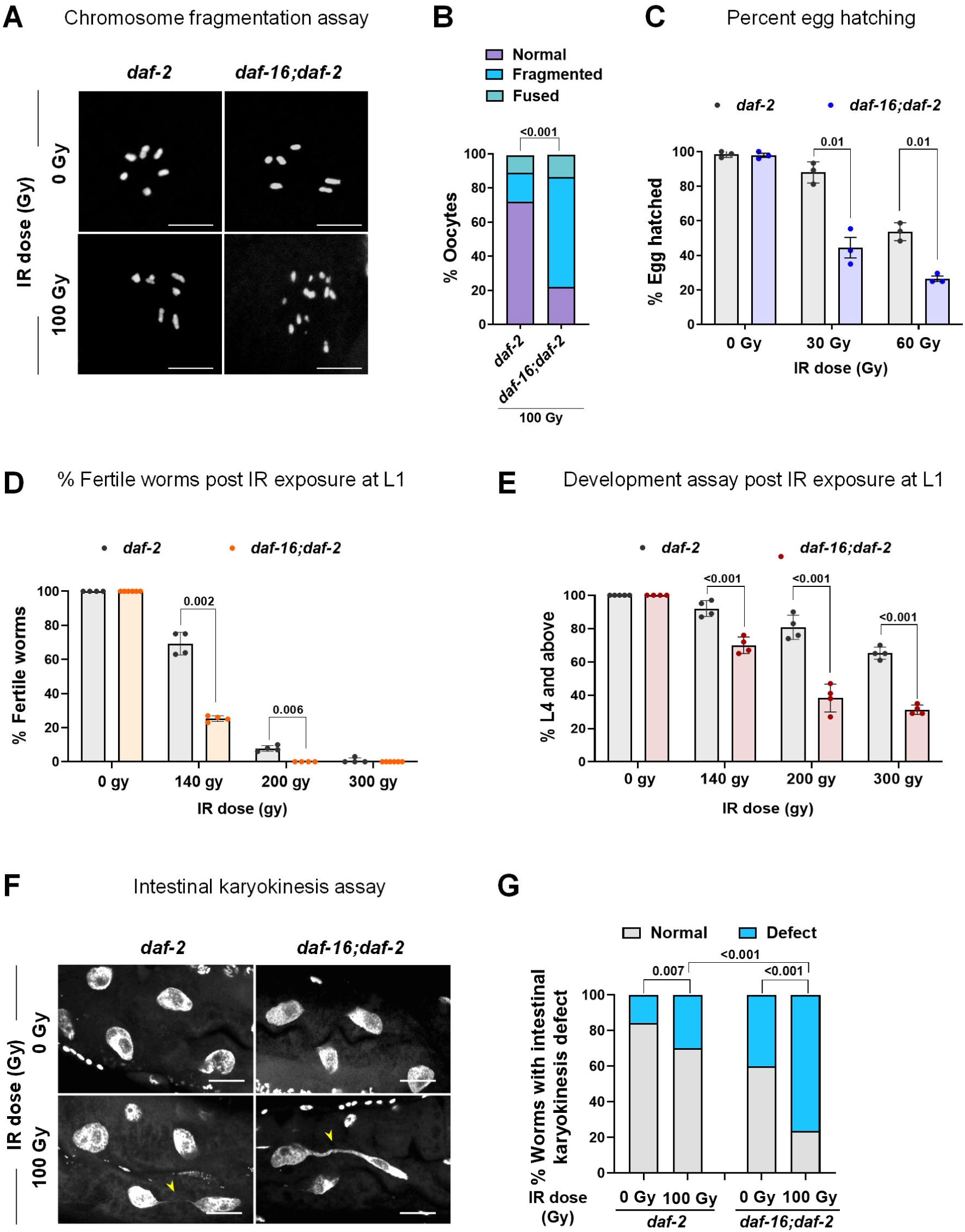
Activated DAF-16 confers increased germline and somatic DDR efficiency following genotoxic stress. **(A)** Representative fluorescence images of DAPI-stained gonads (48 HPI) of *daf-2(e1370)* and *daf-16(mgdf50);daf-2(e1370)* worms grown on control RNAi, irradiated (0 and 100 Gy) at late-L4 stage. Scale bar: 5 µm. **(B)** Quantification for the degree of chromosome fragmentation seen above. The oocyte chromosomes were categorized as normal, fragmented, or fused based on the number of oocyte chromosomes (normal = 6 chromosomes; fragmented = more than 6 chromosomes; fused = less than 6 chromosomes and fusion between two or more chromosomes). Combined data of three biological replicates (*n*≥45) are shown. Statistical comparisons between groups were performed using Chi-square analysis. **(C)** The percentage of hatched eggs of *daf-2(e1370)* and *daf-16(mgdf50);daf-2(e1370)* worms grown on control RNAi and irradiated (0, 30, and 60 Gy) at the late-L4 stage. Egg hatching was monitored for 12-24 HPI. The average of three biological replicates (*n* ≥ 100 per replicate) is shown. Unpaired *t*-test with Welch’s correction. **(D)** The percentage of fertile worms in *daf-2(e1370)* and *daf-16(mgdf50);daf-2(e1370)* irradiated (0, 140, 200, and 300 Gy) at the L1 stage and grown on control RNAi. The percentage of fertile worms (with eggs in the uterus) was subsequently quantified post-IR (when unirradiated worms reached the late day-1 stage). The average of four biological replicates (*n*≥100 for each experiment) is shown. Unpaired *t*-test with Welch’s correction. **(E)** The percentage of larval development of *daf-2(e1370)* and *daf-16(mgdf50);daf-2(e1370)* worms grown on control RNAi and irradiated (0, 140, 200, and 300 Gy) at the L1 stage. The percentage of worms progressing to the L4 stage and beyond (when unirradiated worms reach the early day-1 stage) was subsequently quantified. The average of four biological replicates (*n* ≥ 80 per replicate) is shown. Unpaired *t*-test with Welch’s correction. **(F)** Representative fluorescence images of DAPI-stained intestinal nuclei (48 HPI) of *daf-2(e1370)* and *daf-16(mgdf50);daf-2(e1370)* worms grown on control RNAi and irradiated (0 Gy and 100 Gy) at the late-L4 stage. The arrow points towards fused intestinal cell nuclei (intestinal karyokinesis defect). Scale bars: 10 µm. **(G)** Quantification of the degree of intestinal karyokinesis defect as seen above. The quality was categorized as normal or with defects based on its morphology, where normal indicates distinct intestinal nuclei without any chromatin bridges or intestinal nuclei fusion, and defect indicates intestinal chromatin bridge or fused intestinal nuclei. The combined data of four biological replicates (*n*≥120 intestinal cells) is shown. Statistical comparisons between groups were performed using Chi-square analysis. Experiments were performed at 20°C. Source data are provided in Table 1.

Since proper chromosome segregation at diakinesis is essential for embryonic viability [32], we next assessed egg hatching efficiency after exposing the worms to IR. The IR-induced damage to oocyte chromosomes correlated with a dose-dependent (0, 30, and 60 Gy) reduction in hatching across all strains. However, *daf-16;daf-2* and *daf-16* mutants exhibited significantly lower hatching rates than *daf-2* and WT, respectively (**Fig. 2C, S2E**), and *daf-16;daf-2* mothers produced smaller broods post-IR (**Fig. S2F**). These results underscore the critical role of DAF-16 in maintaining germline genomic stability.

Next, we assessed whether activated DAF-16 in *daf-2* can protect the developing germline from IR-induced damage. At the L1 stage, the worm possesses 2 primordial germline cells and 2 somatic gonad precursor cells, which, during larval development, divide and differentiate to form the complete adult germline [33]. Irradiation (0, 140, 200, and 300 Gy) of worms at the L1 stage resulted in a dose-dependent reduction in fertility. After L1 irradiation, *daf-2* and WT adults retained higher fertility than *daf-16;daf-2* and *daf-16* mutants, respectively (**Fig. 2D, S2G**), suggesting that DAF-16 safeguards germline development under genotoxic stress.

We further explored whether DAF-16 also contributes to somatic DDR. Following L1 irradiation, *daf-16;daf-2* worms exhibited severe developmental delays compared to *daf-2* (**Fig. 2E**), and *daf-16* mutants showed impaired progression to L4 or adulthood relative to WT (**Fig. S2H**). These results indicate that DAF-16 promotes somatic survival and development under DNA damage, consistent with its known role in stress resistance and longevity [34].

To further evaluate somatic DDR, we examined intestinal karyokinesis, a process critical for intestinal cell maturation [35]. Previous studies have linked DNA damage (e.g., IR or oxidative stress) and DDR gene mutations (such as *atm-1*, *dog-1*) to intestinal karyokinesis defects [36] [28] Mirroring these findings, irradiated *daf-16;daf-2* and *daf-16* worms displayed increased intestinal chromatin bridges and binucleation, compared to *daf-2* and WT, respectively (**Fig. 2F, G, S2I, J**), suggesting that amidst genotoxic insults, DAF-16 ensures proper chromosomal segregation during somatic cell division.

Together, these results demonstrate that DAF-16 not only enhances germline DDR but also plays a critical role in somatic genome maintenance, expanding its known functions in stress resistance and aging to include direct modulation of DNA repair mechanisms.

### DAF-16 exerts cell-autonomous and cell non-autonomous control over germline DDR, predominantly through its d/f isoform

DAF-16 exhibits tissue-specific functions to coordinate distinct physiological outcomes. For instance, DAF-16 in germ cells inhibits proliferation, while DAF-16 activation in the hypodermis promotes it [37, 38]. The DAF-16 often performs functions in a cell non-autonomous fashion, as DAF-16 in the proximal gonad has been shown to influence the decline in germline stem cell pool with age [39]. Neuronal and intestinal DAF-16 have been found to be involved in a FOXO-to-FOXO feedback signalling that is required for longevity upon *daf-2* inactivation in either tissue [40]. Interestingly, in a previous study, we elucidated the role of somatic DAF-16 in sensing DNA damage perturbations and influencing the reproductive decision [28]. Therefore, we sought to determine if DAF-16 regulates germline DDR cell-autonomously in the germ cells or cell non-autonomously from the somatic tissues. To achieve germline-specific *daf-16* knockdown, we utilized a specialized RNAi system in which the *rde-1* mutant background (deficient in the Argonaute protein RDE-1) is complemented by reintroducing *rde-1* under a germline-specific *sun-1* promoter [41, 42]. In contrast, somatic knockdown was achieved using a *ppw-1* mutant, which is defective in germline RNAi due to the loss of a PAZ/PIWI-domain protein essential for RNAi efficiency in germ cells [43]. We knocked down *daf-16* in the whole body of *daf-2*, exclusively in somatic tissues (using *daf-2;ppw-1*), or specifically in germ cells of the *daf-2* mutant (using *daf-2;rde-1;sun-1p::rde-1*) by RNAi. We performed the chromosome fragmentation and the egg hatching assays to determine the germline DDR efficiency after control or *daf-16* RNAi in each of these strains. Interestingly, in comparison to control RNAi-fed irradiated worms, we found that *daf-16* knockdown in both germ cells as well as the somatic tissues of the *daf-2* mutants increased chromosomal abnormalities (**Fig. 3A, B**) and dead eggs (**Fig. S3A**) upon irradiation. Notably, the effect on egg hatching was much more pronounced when *daf-16* was knocked-down in the somatic tissues. This suggests that while DAF-16 is essential in both tissue types for implementing the proper DDR mechanism in the germ cells, somatic DAF-16 may play a more crucial role. The dual requirement for DAF-16 in both germline and somatic tissues emphasizes the integrated communication between these compartments, which is essential for maintaining genome stability in the face of genotoxic challenges.

**Fig 3:**
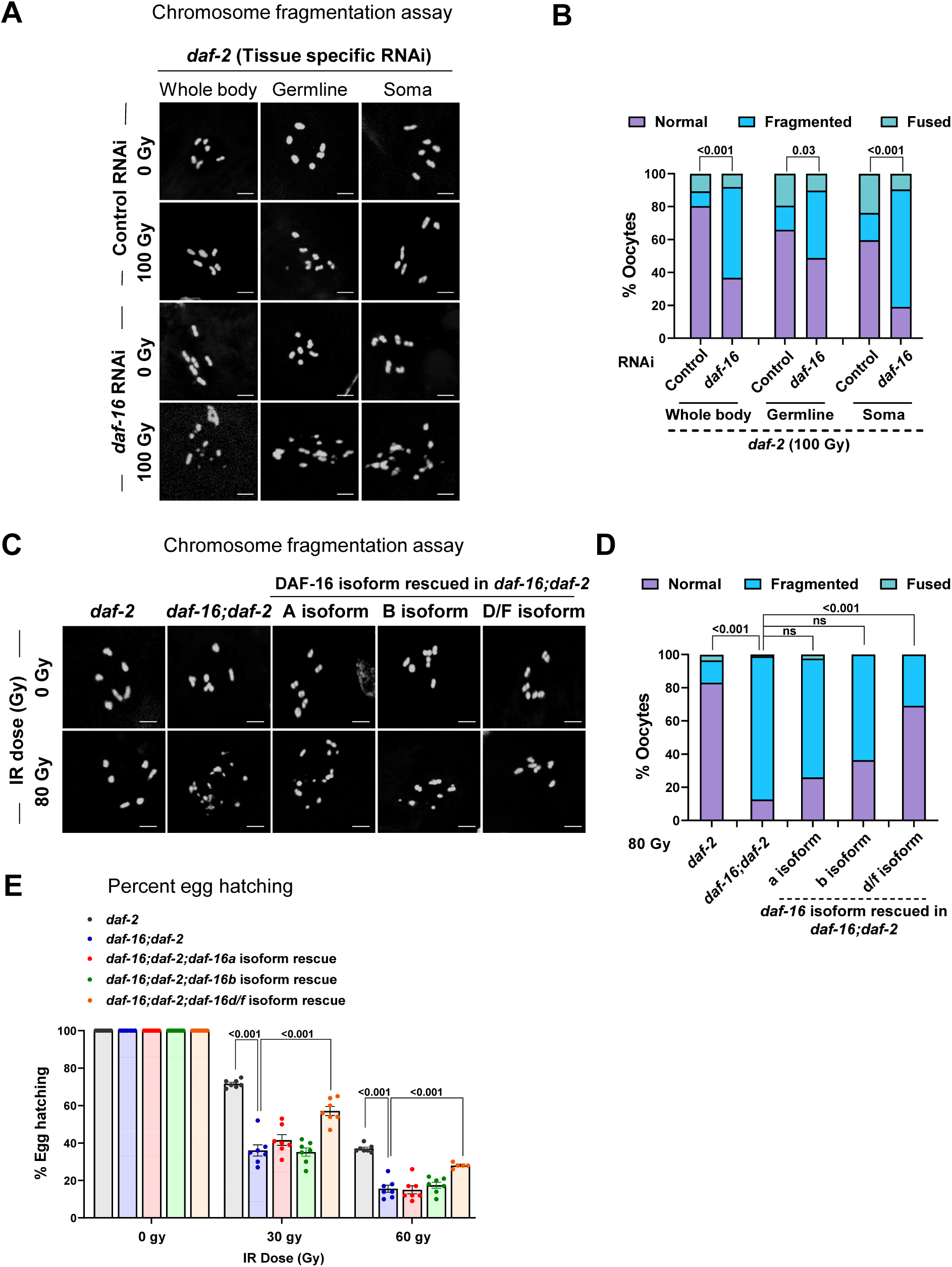
DAF-16 D/F, not A isoform, confers increased germline DDR efficiency following genotoxic stress. **(A)** Representative fluorescence images of DAPI-stained gonads (48 HPI) of *daf-2(e1370)*, *daf-2(e1370)*;*rde-1(mkc36);sun-1p::rde-1* (germline-specific RNAi), and *daf-2(e1370)*;*ppw-1(pk1425)* (soma-specific RNAi) worms grown on control or *daf-16* RNAi, irradiated (0 and 100 Gy) at late-L4 stage. **(B)** Quantification for the degree of chromosome fragmentation as seen above. The oocyte chromosomes were categorized as normal, fragmented, or fused based on their morphology (normal = 6 chromosomes; fragmented = more than 6 chromosomes; fused = less than 6 chromosomes and fusion between two or more chromosomes). Combined data from three biological replicates (*n* ≥ 42) are presented. Statistical comparisons between groups were performed using Chi-square analysis. **(C)** Representative fluorescence images of DAPI-stained gonads (48 HPI) of *daf-2(e1370), daf-16(mgdf50);daf-2(e1370)* and different DAF-16 isoforms rescued *(daf-16a::RFP, daf-16b::CFP, and daf-16d/f::GFP)* in *daf-16(mgdf50);daf-2(e1370)* worms grown on control RNAi, irradiated (0 and 80 Gy) at late-L4 stage. **(D)** Quantification for the degree of chromosome fragmentation as seen above. The oocyte chromosomes were categorized as normal, fragmented, or fused based on their morphology (normal = 6 chromosomes; fragmented = more than 6 chromosomes; fused = less than 6 chromosomes and fusion between two or more chromosomes). Combined data from three biological replicates (*n* ≥ 22) are presented. Statistical comparisons between groups were performed using Chi-square analysis. **(E)** The percentage of hatched eggs of *daf-2(e1370), daf-16(mgdf50);daf-2(e1370)* and different DAF-16 isoforms rescued *(daf-16a::RFP, daf-16b::CFP*, and *daf-16d/f::GFP*) in *daf-16(mgdf50);daf-2(e1370)* worms grown on control RNAi, irradiated (0, 30, and 60 Gy) at the late-L4 stage. Egg hatching was monitored for 12-24 HPI. The average of seven biological replicates (*n* ≥ 80 per replicate) is shown. Two-way ANOVA with Tukey’s multiple comparison test. Experiments were performed at 20°C. Scale bars 5 µm. Source data are provided in Table 1.

The *daf-16* gene codes for multiple isoforms that differ in their promoter regions and transcription start sites. The DAF-16(b) isoform is a smaller protein compared to the DAF-16(a) and DAF-16(d/f) isoforms. The expression pattern is varied amongst these isoforms, with both DAF-16(a) and DAF-16(d/f) expressed more widely than the DAF-16(b) isoform, which is more restricted in expression to the pharynx, a few neurons, and the somatic gonad [8, 44, 45]. The DAF-16(a) isoform predominantly functions to support dauer arrest and longevity of *daf-2*. In the absence of the DAF-16(a) isoform, DAF-16(d/f) plays a role in dauer arrest and longevity [46]. We aimed to investigate which DAF-16 isoform may play a crucial role in DDR. For this, we utilized the different transgenic lines where *daf-16* isoforms (a, b or d/f) are rescued in the *daf-16;daf-2* mutant background. To evaluate the effect on germline DDR, we employed chromosome fragmentation and the egg hatching assays following IR exposure at the late-L4 stage. Interestingly, rescuing the *daf-16(d/f)* isoform in the *daf-16;daf-2* led to a significant alleviation in the IR-induced chromosome fragmentation (**Fig. 3C, D**) and egg hatching (**Fig. 3E**), as compared to the irradiated *daf-16;daf-2* worms. The other *daf-16* isoforms (a or b) did not rescue these phenotypes (**Fig. 3C-E**). The level of chromosome fragmentation and egg hatching percentage was comparable in irradiated *daf-16(d/f)* isoform rescued lines and *daf-2* (**Fig. 3C-E**), indicating a predominant role of DAF-16(d/f) isoform in the germline DDR in the *daf-2* worms.

Further, we wanted to assess the contribution of each of the *daf-16* isoforms in somatic DDR, for which we employed the L1 larval IR sensitivity assay (development and fertility). However, none of the isoforms could rescue the developmental defects (**Fig. S3B**) or the sterility (**Fig. S3C**) of the worms that were irradiated at L1.

These findings underscore the specific importance of the *daf-16(d/f)* isoform in maintaining germline DDR efficiency. It further suggests that somatic DDR may either require a distinct *daf-16* isoform or involve a combinatorial action of multiple isoforms to fully preserve DNA repair capacity following genotoxic stress.

### DAF-16 mediates higher DDR efficiency in *daf-2* following genotoxic stress

We observed significantly higher frequencies of oocyte chromosomal defects (fragmentation and fusion) in *daf-16;daf-2* compared to *daf-2* mutants at 48 hours post-IR (100 Gy at late-L4 stage; **Fig. 2A,B**). This DAF-16-dependent phenotype suggested either differential initial damage or repair efficiency between the strains. To distinguish these possibilities, we performed a comprehensive time-course analysis of chromosomal integrity at 12, 18, 24, 30, 36, 42, and 48 HPI. Intriguingly, while *daf-2* oocytes consistently maintained low levels of chromosomal abnormalities throughout all time points (**Fig. 4A, S4A, B**), *daf-16;daf-2* oocytes exhibited progressive accumulation of damage (**Fig. 4A, S4A, B).** Notably, the difference in chromosomal abnormalities between *daf-2* and *daf-16;daf-2* was minimal (19.2%) at 12 HPI, but became increasingly significant at later time points (72.65 at 18-48 hours) (**Fig. 4A S4A, B)**. This temporal pattern suggests that, while both strains incur almost similar initial damage, activated DAF-16 may drive the subsequent repair processes efficiently in *daf-2*. The progressive divergence in chromosomal integrity phenotypes supports a model where DAF-16 activity sustains DNA repair capacity over time, rather than simply preventing initial damage, consistent with a conserved role of FOXO3a in double-strand break repair [47]

**Fig 4:**
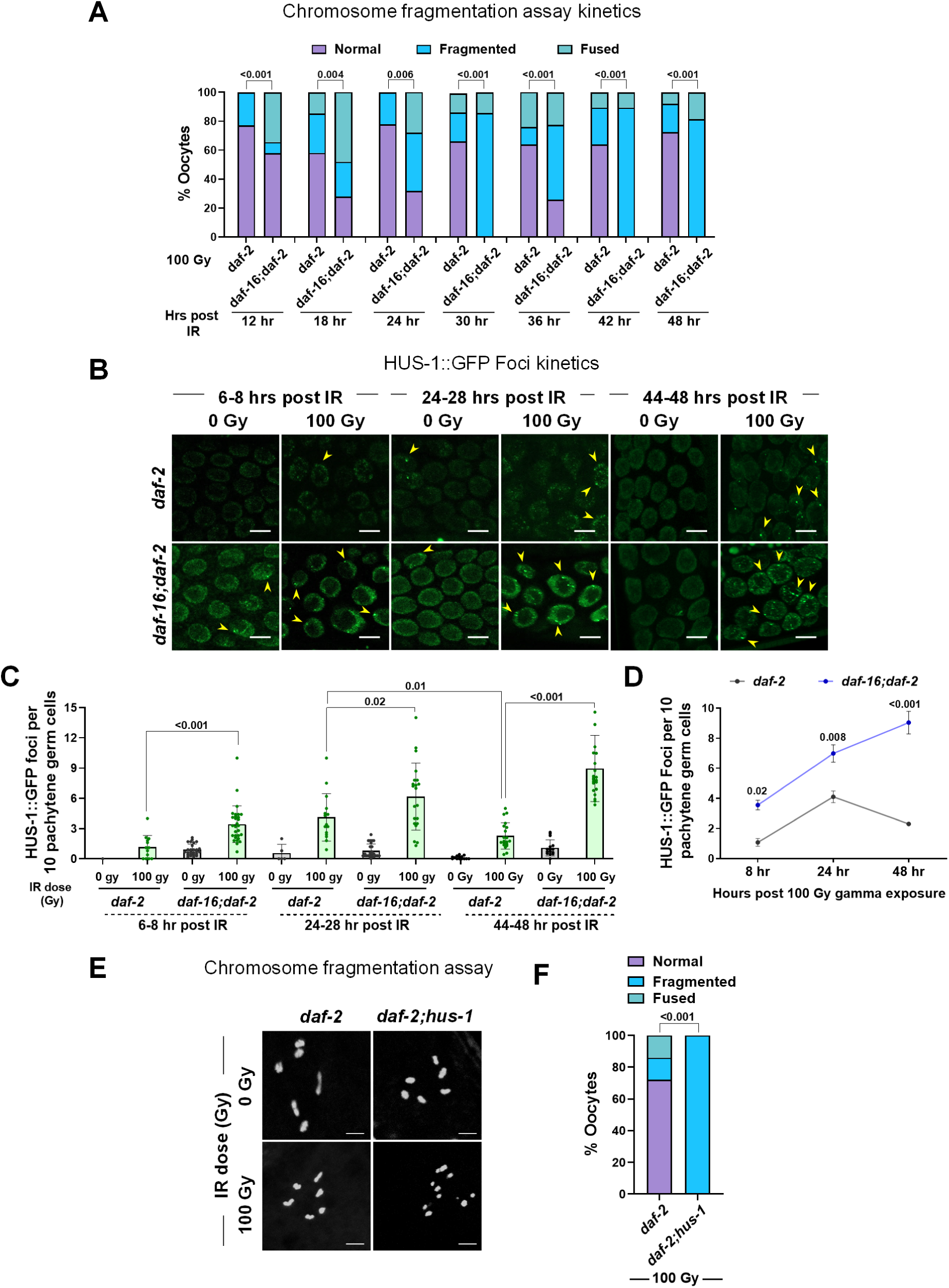
Kinetics of DNA damage repair following genotoxic stress is accelerated upon DAF-16 activation. **(A)** Quantification of the chromosomal fragmentation kinetics of DAPI-stained gonads (12, 18, 24, 30, 36, 42, and 48 HPI) in *daf-2(e1370)* and *daf-16(mgdf50);daf-2(e1370)* worms grown on control RNAi, irradiated (100 Gy) at the late-L4 stage (as seen in Figure S4A,B). The oocyte chromosomes were categorized as normal, fragmented, or fused based on their morphology (normal = 6 chromosomes; fragmented = more than 6 chromosomes; fused = less than 6 chromosomes and fusion between two or more chromosomes). Combined data from three biological replicates (*n* ≥ 22) are presented. Statistical comparisons between groups were performed using Chi-square analysis. **(B)** Representative fluorescence images showing HUS-1::GFP (DNA damage checkpoint protein) expression in *daf-2(e1370)* and *daf-2(e1370);daf-16(mgdf50)* (6-8, 24-28 and 44-48 HPI). Worms grown on control RNAi were irradiated (0 and 100 Gy) at the late-L4 stage. Yellow arrows pointed towards HUS-1::GFP foci formation. **(C)** Quantification for the HUS-1::GFP foci per 10 pachytene germ cells per gonad as seen above. Combined data from three biological replicates (*n* ≥ 12) are presented. Unpaired *t*-test with Welch’s correction. **(D)** Quantification for the HUS-1::GFP foci per 10 pachytene germ cells per gonad as seen in Fig. 4B. Combined data from three biological replicates (*n* ≥ 12) are presented. Two-way ANOVA with Tukey’s multiple comparison test. **(E)** Representative fluorescent images of DAPI-stained gonads (48 HPI) of *daf-2(e1370)* and *daf-2(e1370);hus-1(op241)* worms grown on control RNAi, irradiated (0 and 100 Gy) at the late-L4 stage. **(F)** Quantification for the degree of chromosome fragmentation as seen above. The oocyte chromosomes were categorized as normal, fragmented, or fused based on its morphology (normal = 6 chromosomes; fragmented = more than 6 chromosomes; fused = less than 6 chromosomes and fusion between two or more chromosomes). Combined data from three biological replicates (*n* ≥ 48) are presented. Statistical comparisons between groups were performed using Chi-square analysis Experiments were performed at 20°C. Scale bars 5 µm. Source data are provided in Table 1.

To further assess the role of FOXO in the damage repair efficiency, we examined the localization of the checkpoint protein HUS-1::GFP in germ cells following IR. We utilized the transgenic strain *hus-1(op241);hus-1p::hus-1::gfp* to visualize HUS-1 dynamics following IR. In untreated worm gonads, HUS-1 remains diffused in the cytoplasm, with only a few HUS-1 foci detected in the germ cell nuclei [48] However, upon IR treatment, HUS-1 accumulates at the sites of DNA damage, leading to the formation of HUS-1 foci, which are known to facilitate the recruitment of other DNA damage repair proteins [48, 49]. Upon IR exposure, we observed an increase in HUS-1::GFP foci formation (8 hrs and 27 hrs post-IR) in both *daf-2* and *daf-16;daf-16* pachytene germ cells. However, the extent of HUS-1::GFP foci formation was markedly elevated in the *daf-16;daf-2* gonads in comparison to the *daf-2* worms (**Fig. 4B, C, D**), suggesting that the absence of DAF-16 exacerbates DNA damage accumulation. Moreover, at 48 hrs post-IR, while the number of HUS-1::GFP foci declined in the irradiated *daf-2* germ cells, HUS-1::GFP foci were still increased in the *daf-16;daf-2* germ cells (**Fig. 4B, C, D**). This, along with DAF-16-dependent lower chromosomal abnormalities in *daf-2* gonads following irradiation, highlights the role of DAF-16 in conferring increased DDR efficiency under low insulin signalling.

To further investigate the role of HUS-1 in *daf-2* DDR in the germline, we analysed *daf-2;hus-1* double mutants following IR. Reduction of HUS-1 function in *daf-2* led to a massive increase in chromosome fragmentation following IR (**Fig. 4E, F**). This highlights the crucial role of the conserved DNA damage checkpoint protein, HUS-1, in maintaining higher DDR efficiency under conditions of low insulin signalling.

### DAF-16 employs canonical DDR, but not apoptosis, to enhance repair efficiency

Upon exposure to a DNA-damaging agent, the DDR machinery in the germline is activated to repair the damaged DNA. If the damage remains unrepaired, germ cell apoptosis is triggered to eliminate the damaged/defective germ cells [1]. We speculated that the germ cell apoptosis in *daf-2* gonads might also be elevated to support enhanced clearance of damaged nuclei. To monitor the kinetics for germ cell apoptosis in *daf-2* and *daf-16;daf-2*, we utilized the transgenic strain *ced-1p::gfp*, where CED-1 is a transmembrane protein on the surface of sheath cells that engulf the dying germ cells [50]. At 6 and 12 HPI, an increased number of apoptotic corpses was observed in both *daf-2* and *daf-16;daf-2* gonads. However, at 24 and 36 HPI, apoptosis was attenuated in irradiated *daf-2* germlines, whereas a significant increase in germ cell apoptosis persisted in the irradiated *daf-16;daf-2* gonads (**Fig. 5A, B, S5A, B**). This contrasted with our expectation of increased apoptotic clearance of germ cells in irradiated *daf-2* germline. However, a faster kinetics of apoptosis in *daf-2* may hint towards a lower burden of damaged germ cells due to an enhanced damage repair efficiency. To further explore the role of apoptosis, we examined *daf-2;ced-4* [CED-4 is the *C. elegans* ortholog for mammalian APAF1 that activates the procaspase-9 [51]]. Despite the absence of apoptotic clearance, *daf-2;ced-4* worms did not exhibit increased chromosomal abnormalities compared to *daf-2* alone (**Fig. 5C, D**). Moreover, knock-down of the worm caspase, CED-3 [52] by RNAi also did not exacerbate the chromosomal defects in *daf-2* mutant worms upon irradiation (100Gy) (**Fig. S5C, D**). These results suggests that the enhanced DDR in *daf-2* is not mediated by apoptosis but rather by improved DNA repair.

**Fig 5:**
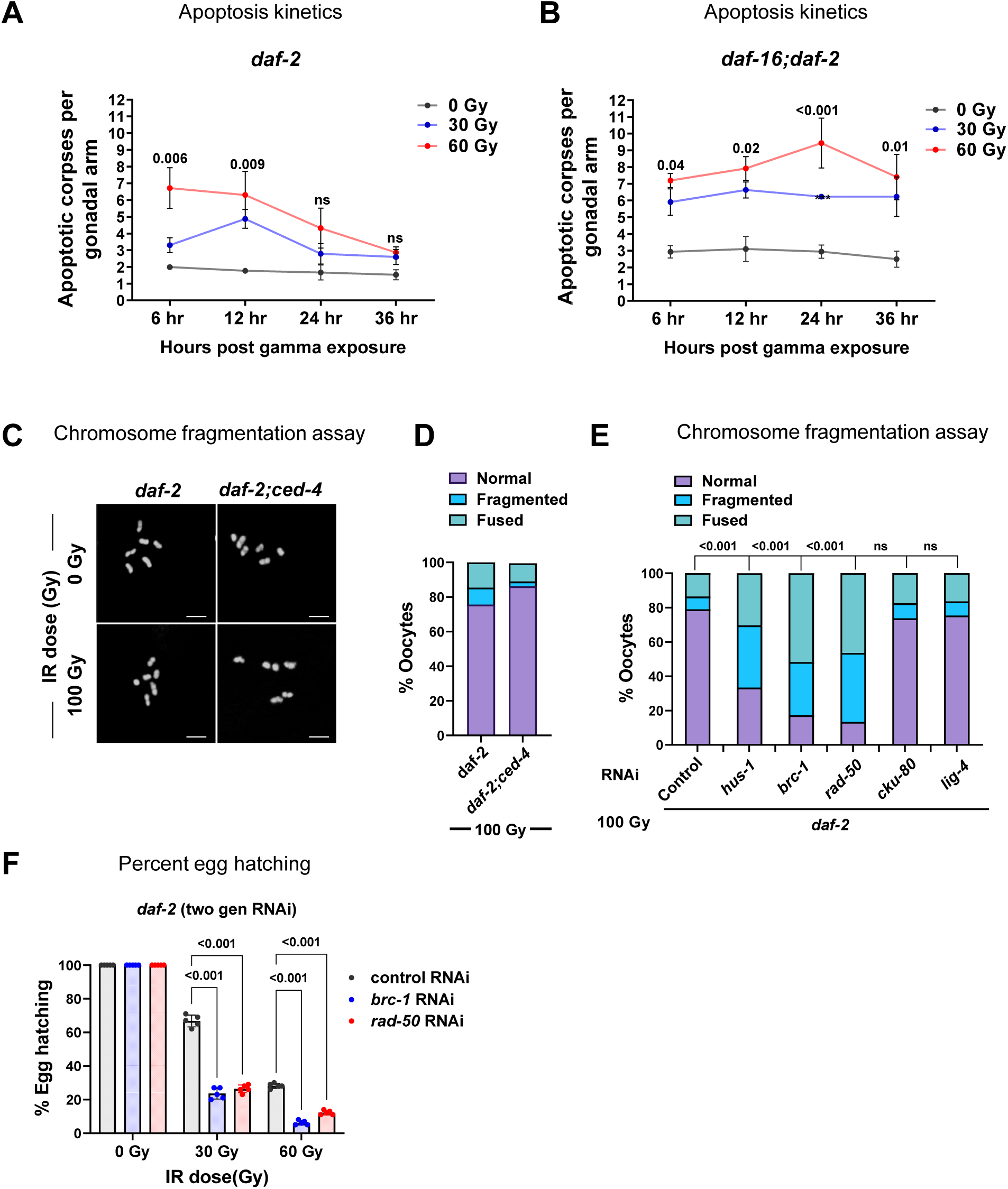
DAF-16 requires canonical DDR components, and not apoptosis, to enhance DDR efficiency. **(A)** Kinetics of apoptosis (6, 12, 24, and 36 HPI) showing mean apoptotic corpse numbers per gonadal arm of *daf-2(e1370);ced-1::GFP* worms grown on control RNAi, irradiated (0, 30, and 60 Gy) at the late-L4 stage (as seen in Figure S5A). The average of three biological replicates is shown (n≥17 for each replicate). Two-way ANOVA with Tukey’s multiple comparison test. **(B)** Kinetics of apoptosis (6, 12, 24, and 36 HPI) showing mean apoptotic corpse numbers per gonadal arm of *daf-16(mgdf50);daf-2(e1370);ced-1::GFP* worms grown on control RNAi, irradiated (0, 30, and 60 Gy) at late-L4 stage (as seen in Figure S5B). An average of three biological replicates (n≥22 for each replicate). Two-way ANOVA with Tukey’s multiple comparison test. **(C)** Representative fluorescence images of DAPI-stained gonads (48 HPI) of *daf-2(e1370)* and *daf-2(e1370);ced-4(n1162)* worms grown on control RNAi, irradiated (0 and 100 Gy) at the late-L4 stage. Scale bars: 5 μm. **(D)** The quantification of the degree of chromosome fragmentation as seen above. The oocyte chromosomes were categorized as normal, fragmented, or fused based on their morphology (normal = 6 chromosomes; fragmented = more than 6 chromosomes; fused = less than 6 chromosomes and fusion between two or more chromosomes). Combined data from three biological replicates (*n* ≥ 36) are presented. Statistical comparisons between groups were performed using Chi-square analysis. **(E)** Quantification of the extent of chromosomal fragmentation of DAPI-stained gonads (48 HPI) in *daf-2(e1370)* worms grown for two generations on control, *hus-1*, *brc-1*, *rad-50*, *cku-80*, and *lig-4* RNAi, irradiated (0 and 100 Gy) at the late-L4 stage (as seen in Figure S5E). Based on morphology, the oocyte chromosomes were categorized as normal, fragmented, or fused (normal = 6 chromosomes; fragmented = more than 6 chromosomes; fused = less than 6 chromosomes and fusion between two or more chromosomes). Combined data from four biological replicates (*n* ≥ 58) are presented. Statistical comparisons between groups were performed using Chi-square analysis. **(F)** The percentage of hatched eggs of *daf-2(e1370)* worms grown for two generations on control, *brc-1*, and *rad-50* RNAi, irradiated (0, 30, and 60 Gy) at the late-L4 stage. Egg hatching was monitored for 12-24 HPI. Average of five biological replicates (*n* ≥ 120 per replicate). Two-way ANOVA with Tukey’s multiple comparison test. Experiments were performed at 20°C. Source data are provided in Table 1.

As HUS-1 is required to maintain germline DDR in *daf-2* (**Fig. 4D, E**), we next wanted to ascertain the requirement of other key proteins involved in the canonical DDR. For this, we exposed *daf-2* worms to RNAi-expressing bacteria, followed by IR exposure at the late-L4 stage, and examined chromosome fragmentation in the oocytes. We were able to validate the functionality of our KD experiments since *hus-1* KD by RNAi led to a significant increase in chromosome fragmentation (**Fig. 5E, S5E**) post-IR, similar to that observed in the irradiated *daf-2;hus-1* mutant (**Fig. 4E, F**). Also, *rad-51* KD led to chromosomal abnormalities even in the unirradiated *daf-2* worms (**Fig. S5E**). Compared to worms grown on control RNAi, *daf-2* mutants subjected to knockdown of DDR genes (*hus-1, brc-1* or *rad-50*) exhibited significantly higher levels of chromosomal abnormalities (**Fig. 5E, S5E**) and reduced egg hatchability (**Fig. 5F**) following irradiation. Notably, homologous recombination (HR)-mediated DNA repair, which depends on components such as HUS-1, BRC-1, and RAD-50, appears to be essential for maintaining genome integrity under these conditions. In contrast, non-homologous end joining (NHEJ) was found to be largely dispensable, as RNAi-mediated depletion of *cku-80* and *lig-4* [cku-80 is part of the double-strand breaks (DSBs) recognizing CKU70/CKU80 heterodimer and lig-4 is the NHEJ ligase [31]] did not lead to a detectable increase in chromosomal defects (**Fig. 5E, S5E**). This is consistent with previous findings showing that HR is the predominant repair pathway for DSBs in the germline, whereas post-mitotic somatic cells primarily rely on NHEJ for DSB repair [31]

Importantly, these findings indicate that the enhanced DNA damage response observed due to activation of DAF-16 (in *daf-2* mutants) is not due to increased apoptotic clearance of damaged germ cells post-IR, but rather reflects an improved capacity for repair, mediated by the robust function of the core DDR machinery.

## Discussion

In the current study, we uncover a previously underexplored role of the longevity-linked IIS axis in the DDR. Using *Caenorhabditis elegans* long-lived Insulin-IGF-1 receptor mutant (*daf-2*), we show up-regulation in the basal levels of DDR genes due to binding of activated worm FOXO transcription factor, DAF-16, to DDR gene promoters. We used IR to elicit DNA damage and find a DAF-16-dependent increase in the DNA-repair capacities in both mitotically/meiotically active (germline cells) and post-mitotic cells (somatic cells). Our findings reinforce an evolutionarily well-conserved role of FOXO as custodian of the genome that helps maintain DNA integrity (**Fig. 6**).

**Fig 6.**
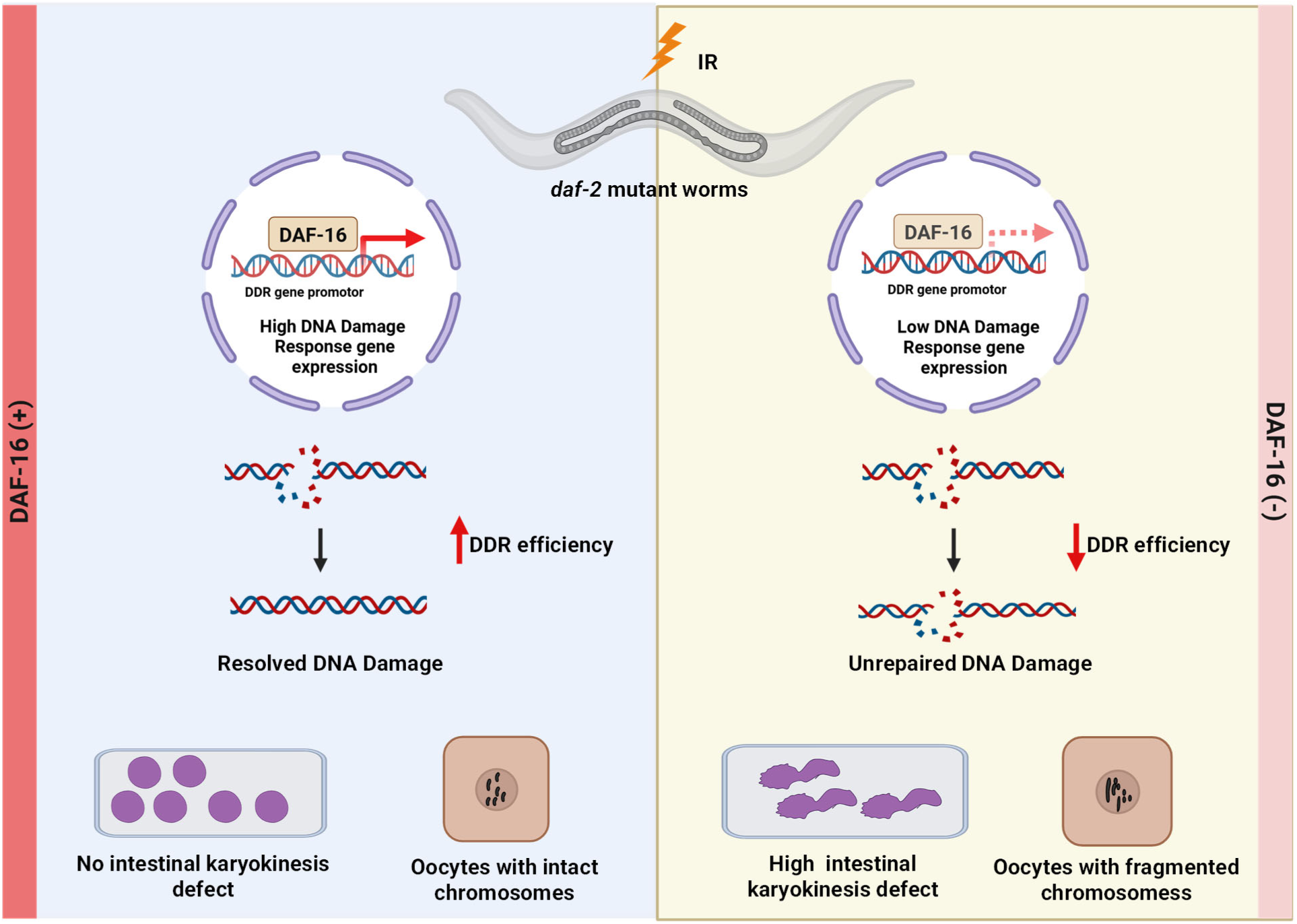
Model depicting the role of DAF-16 in somatic and germline DDR. Upon genotoxic stress (such as IR: Ionizing radiation), DNA damage such as double-strand break is elicited. In *daf-2* mutant worms (characterised by low insulin signalling), the presence of the activated DAF-16/FOXO transcription factor results in an increased level of DNA damage response (DDR) genes. This leads to a heightened DDR efficiency and genomic stability in both somatic and germ cells. However, the absence of DAF-16 in *daf-2* mutant animals compromises DDR efficiency, leading to unresolved DNA damage that manifests as karyokinesis defects in intestinal cells and as fragmented chromosomes in oocytes. Thus, DAF-16 safeguards genome integrity in both somatic and germ cells. The Schematic was created using BioRender.com.

Previously, in mammalian cell line models, FOXO has been shown to protect against DNA damage by inducing cell-cycle arrest [11], thus allowing time for DNA damage to be repaired [53] or by triggering cell cycle exit or apoptosis in case of unresolved DNA damage [14, 16, 54, 55]. Overexpression of FOXO3a in primary human fibroblasts has been shown to decrease DSB-induced γH2AX foci and suppress genomic rearrangements [47]. FOXO3 has also been shown to protect the hematopoietic stem and progenitor cells from oxidative stress-induced DNA damage [13]. We wanted to test whether and how the worm FOXO transcription factor, DAF-16, plays a functional role in DNA damage repair. For examining germ-line DDR, we observed oocyte chromosome fragmentation and egg hatching following IR. In germline cells, *daf-2* mutants maintained intact bivalent chromosome structures post-IR exposure, in contrast to wild-type (WT) or *daf-16;daf-2* double mutants, which displayed significant chromosomal fragmentation. This DAF-16-dependent protection also translated into improved embryonic viability following IR-induced stress. These results underscore a functional role for DAF-16 in promoting germline genome stability. While the germline DDR is relatively well characterized, the understanding of somatic DDR in *C. elegans* is limited and partially repressed by the DREAM complex [56, 57]. We evaluated somatic repair by quantifying IR-induced developmental arrest and intestinal karyokinesis defects [36]. We found increased larval development post-IR in *daf-2* compared to the *daf-16;daf-2* mutant. In many DDR mutants (*atm-1, dog-1*) or on treating animals with chemicals that generate reactive oxygen species (ROS), the formation of intestinal chromatin bridges or fusion of intestinal nuclei have been noticed [36]. Utilizing the intestinal karyokinesis defects post-IR as a readout to assess somatic DDR efficiency, we found fewer karyokinesis defects in irradiated *daf-2* mutants than in *daf-16;daf-2* worms. Interestingly, our findings contrast with prior studies reporting that DAF-16 promotes larval development despite unresolved UV-induced DNA lesions [58], suggesting that under IR stress, DAF-16 enhances DNA repair rather than bypassing damage. Even in the WT background, where DAF-16 is predominantly cytoplasmic, removal of the transcription factor compromised somatic and germline DDR capabilities.

Since germline maintenance is an energetically demanding process, the somatic tissues often take inputs from the environment to influence germline development [59], proliferation [60], apoptosis [61], oocyte growth and maturation [62]. Increasing evidence supports the regulatory role of soma in germline DDR. For instance, the somatic niche cells have been shown to influence CEP-1/p53-mediated DDR in primordial germ cells (PGC) and mammalian stem cells [63]. Similarly, the stress-responsive PMK-1/p38 functions in the intestine to promote stress-induced (heat, IR) apoptosis in germline [64]. Neuronal HIF-1, IRE-1, and intestinal KRI-1 function upstream or in parallel to CEP-1/p53 to regulate IR-induced germ cell apoptosis [65–67]. Quite strikingly, in our study, we found knocking down *daf-16*, either in the soma or the germline, led to increased chromosomal defects and embryonic lethality post-IR exposure. Strikingly, these effects were more contrasting upon somatic DAF-16 depletion. These findings establish that while DAF-16 functions both cell-autonomously in germ cells and cell non-autonomously in the somatic cells to influence germline DDR, somatic DAF-16 appears to play a predominant role in germline DDR repair. Additionally, using isoform-specific rescue lines, we show DAF-16 (d/f) isoform to be crucial for germline DDR, whereas somatic DDR appears to require the combined activity of multiple DAF-16 isoforms, suggesting a tissue-specific complexity in isoform function. In the context of lifespan regulation, the neuronal and the intestinal DAF-16 have been shown to form a feedback loop, and both are required for longevity induced by neuronal or intestinal IIS reduction [40]. It is therefore plausible that similar inter-tissue communication underlies the DAF-16-mediated coordination of DDR. Whether this cross-talk is mediated by insulin-like peptides, neuropeptides, hormonal signals, or other small molecules remains to be explored.

To monitor the DDR kinetics, we utilized HUS-1::GFP (a checkpoint protein fused to gfp [48]) foci detection, chromosomal abnormalities, and apoptosis kinetics following IR exposure. Through the chromosome fragmentation kinetics, we observed that, while both strains incur almost similar initial damage, activated DAF-16 may drive the subsequent repair processes efficiently in *daf-2*. This indicates an enhanced repair capacity owing to heightened DAF-16 activity in *daf-2,* rather than simply preventing initial damage. Lower levels of HUS-1::GFP foci formation and accelerated resolution of IR-induced apoptosis in *daf-2* support these findings. Interestingly, the canonical components of the HR arm of DSB repair, such as *hus-1*, *brc-1*, and *rad-50*, were found to be critical for robust germline DDR, while the NHEJ (*cku-80* and *lig-4*) arm of DSB repair [31] or apoptotic machinery were found to be dispensable. These results suggest that DAF-16 may selectively promote HR-mediated repair over alternative pathways during genotoxic stress in germ cells, where the preservation of genomic integrity is critical for reproductive success and organismal fitness. Previously, FOXO3 has been shown to physically interact with the Ataxia-Telangiectasia Mutated (ATM: a protein kinase and a master regulator of DDR [17] and promote intra-S-phase or G2/M cell-cycle checkpoint, and repair of damaged DNA [68]. In line with this, it would be worthwhile to test if, apart from transcriptional regulation of DDR genes, DAF-16 also physically interacts with the DDR components to enhance damage recognition and resolution, thereby orchestrating a rapid and effective DNA damage response.

With its role in cell cycle arrest [69, 70] and apoptosis [54, 71, 72], FOXO is generally regarded as a tumor suppressor [73–75] However, emerging studies complicate this narrative [76–78]. In cancers such as neuroblastoma, colon and thyroid carcinoma, FOXO activity has been linked to poor prognosis [77, 79, 80]. As an extension of our current findings, we speculate that FOXO activation may enhance the DDR efficiency in cancer cells and may cause resistance against chemotherapeutic drugs which normally induce cancer cell death by causing DNA damage. Therefore, while FOXO activation may be beneficial in normal cells for genome maintenance, its role in cancer must be carefully contextualized to avoid unintended therapeutic resistance.

## Materials and Methods

### *C. elegans* strain maintenance

Unless otherwise mentioned, all the *C. elegans* strains were maintained and propagated at 20°C on *E. coli* OP50 using standard procedures [81]. The strains used in this study are provided in Table 2.

### Preparation of RNAi plates

RNAi plates were prepared using an autoclaved nematode growth medium supplemented with 100 μg/ml ampicillin and 2 mM IPTG. The plates were allowed to dry at room temperature for 24 hours. Bacterial cultures containing the specific RNAi construct were grown overnight in Luria-Bertani (LB) medium supplemented with 100 μg/ml ampicillin and 12.5 μg/ml tetracycline at 37°C in a shaker incubator. The following day, the saturated primary culture was diluted 1:50 in fresh LB medium containing 100 μg/ml ampicillin and incubated at 37°C in a shaker until the optical density at 600 nm (OD₆₀₀) reached 0.5–0.6. Bacterial cells were then pelleted by centrifugation at 3214 × *g* for 5 minutes at 4°C and resuspended in 1/10th of the original volume in M9 buffer containing 100 μg/ml ampicillin and 1 mM IPTG. Approximately 350 μl of this bacterial suspension was seeded onto RNAi plates and allowed to dry at room temperature for 48 hours before being stored at 4°C until further use.

### Hypochlorite treatment for synchronizing worm population

Gravid adult worms, initially grown on *E*. *coli* OP50 bacteria, were collected using M9 buffer in a 15 ml falcon tube. Worms were washed twice by first centrifuging at 652 g for 60 seconds followed by resuspension of the worm pellet in 1X M9 buffer. After the second wash, the worm pellet was resuspended in 3.5 ml of 1X M9 buffer and 0.5 ml 5N NaOH and 1 ml of 4% Sodium hypochlorite solution was added. The mixture was vortexed for 4–5 minutes until the entire worm body dissolved, leaving behind the eggs. The eggs were washed 5–6 times, by first centrifuging at 1258 g, the 1X M9 was aspirated out, followed by resuspension in a fresh 1X M9 buffer to remove traces of bleach and alkali. After the final wash, eggs were kept in 15 ml falcons with ∼ 10 ml of 1X M9 buffer and kept on rotation ∼15 r.p.m for 17 hours to obtain L1 synchronized worms for all strains. The L1 worms were obtained by centrifugation at 805 g followed by resuspension in approximately 200–300 μl of M9 and added to respective RNAi plates.

### RNA isolation

L4-staged worms were collected using 1X M9 buffer and washed thrice to remove bacteria. Trizol reagent (200 μl; Takara Bio, Kusatsu, Shiga, Japan) was added to the 50 μl worm pellet and subjected to three freeze-thaw cycles in liquid nitrogen with intermittent vortexing for 1 minute to break open the worm bodies. The samples were then frozen in liquid nitrogen and stored at -80°C till further use. Later, 200 μl of Trizol was again added to the worm pellet and the sample was vigorously vortexed for 1 minute. To this, 200 μl of chloroform was added and the tube was gently inverted several times followed by 5 minutes of incubation at room temperature. The sample was then centrifuged at 12000 g for 15 minutes at 4°C. The RNA containing the upper aqueous phase was gently removed into a fresh tube without disturbing the bottom layer and interphase. To this aqueous solution, an equal volume of isopropanol was added and the reaction was allowed to sit for 10 minutes at room temperature followed by centrifugation at 12000 g for 15 minutes at 4°C. The supernatant was carefully discarded without disturbing the RNA-containing pellet. The pellet was washed using 1 ml 70% ethanol solution followed by centrifugation at 8000g for 10 minutes at 4°C. The RNA pellet was further dried at room temperature and later dissolved in autoclaved MilliQ water followed by incubation at 65°C for 10 minutes with intermittent tapping. The concentration of RNA was determined by measuring absorbance at 260 nm using a NanoDrop UV spectrophotometer (Thermo Scientific, Waltham, USA) and the quality was checked using denaturing formaldehyde-agarose gel.

### Gene expression analysis using quantitative real-time PCR (QRT-PCR)

First-strand cDNA synthesis was carried out using the Iscript cDNA synthesis kit (Biorad, Hercules, USA) following the manufacturer’s guidelines. The prepared cDNA was stored at -20°C. Gene expression levels were determined using the Brilliant III Ultra-Fast SYBR Green QPCR master mix (Agilent, Santa Clara, USA) and Agilent AriaMx Real-Time PCR system (Agilent, Santa Clara, USA), according to manufacturer’s guidelines. The relative expression of each gene was determined by normalizing the data to *actin* expression levels. The list of primers is provided in Table 3.

### Measurement of cell corpses using CED-1::GFP

Engulfed cell corpses were analyzed using transgenic *C. elegans* expressing CED-1::GFP, a transmembrane protein localized to surrounding sheath cells responsible for engulfing apoptotic cell corpses. *daf-2(e1370);ced-1p::ced-1::GFP* (*smIs34*) and *daf-16(mgdf50);daf-2(e1370);ced-1p::ced-1::GFP* (*smIs34*) worms were bleached, and their eggs were incubated in a 1X M9 buffer for 17 hours to obtain synchronized L1 larvae. Approximately 100 L1 worms were then transferred onto control RNAi plates in triplicate. The worms were irradiated (0, 30 and 60 gy) at the late-L4 stage and imaged (6, 12, 24 and 36 HPI) in Z-stacks using a 488 nm laser to excite GFP on an LSM980 confocal microscope (Carl Zeiss, Oberkochen, Germany). The images were processed into maximum intensity projections (MIPs), and the number of cell corpses per gonadal arm was manually quantified.

### Chromosomal fragmentation assay

Approximately 100 L1 worms were placed onto control or respective RNAi plates in triplicate. The worms were irradiated with ionizing radiation (IR) of different doses at the L4 stage. At the desired time point post-irradiation, the worms were stained with DAPI, and the oocyte chromosomes were imaged in *z*-stack using a using a 405 nm laser to LSM980 confocal microscope (Zeiss). For scoring chromosome fragmentation, images were converted into maximum intensity projections (MIPs) and scored. The oocyte chromosomes were categorized as normal, fragmented, or fused based on its morphology. Normal oocytes contain six bivalent chromosomes. Fragmented oocytes exhibit more than six and clustered chromosomes, while fused oocytes appear as aggregates with fewer than six bivalent chromosomes.

### DAPI staining

Worms were cultured on specific RNAi plates from the L1 stage onward. Adult worms were collected in a 1.5 ml Eppendorf tube containing 1X M9 buffer and allowed to settle. The buffer was carefully removed using a glass Pasteur pipette, leaving approximately 100 μl of worm suspension. Ice-cold 100% methanol (1 ml) was then added to the worm pellet, followed by incubation at -20°C for 30 minutes. The methanol-fixed worm pellet was placed on a glass slide, and after the methanol had evaporated, Fluoroshield with DAPI (Invitrogen, Carlsbad, USA) was applied for staining. A coverslip was carefully placed on top, ensuring no air bubbles, and sealed with transparent nail polish. Finally, images were acquired using a 405 nm laser excitation for DAPI on an LSM980 confocal microscope (Carl Zeiss, Oberkochen, Germany).

### L1 larval development and fertility assay

Almost 100 L1 worms were irradiated with different dose of gamma (0, 140, 200 and 300 Gy) and scored for worms progressing to the L4-stage or beyond (when the unirradiated worms reached early Day-1 adult). The larval development was assessed by determining the percentage of developed larvae that reached L4 stage or above. In the same experiment, fertile worms were counted (with eggs in the uterus) when the unirradiated controls reached late-day1 of adulthood. The percentage fertile worms for each condition were then calculated.

### Egg hatching

To assess the extent of DNA damage repair in the germline the percentage of hatched eggs was determined. Worms were grown on respective RNAi L1 onwards, and gamma irradiated (0, 30 and 60 gy) at the late-L4 stage. Almost 15 adult worms per condition were allowed to lay eggs for 12-24 hr window post IR. The mother worms were then sacrificed and the number of eggs on respective plates were counted. After 48 hours the number of hatched progenies were counted, and the percentage of hatched eggs were calculated for each condition.

### Brood size

Worms were grown on control RNAi from L1 onwards and gamma irradiated (0, 30 and 60 gy) at the late-L4 stage. Upon reaching the young adult stage, five worms were picked onto fresh RNAi plates, in triplicates, and allowed to lay eggs for 24 hours. The worms were then transferred to fresh plates every day until worms ceased to lay eggs, and these plates were counted after 48 hours to document the number of hatched worms. The total number of hatched progenies per worm is defined as brood size.

### Intestinal karyokinesis

Approximately 100 L1 worms were placed onto control RNAi plates in triplicate. The worms were irradiated (0 and 100 Gy) at the L4 stage. At 48 HPI, the worms were stained with DAPI, and the intestinal nuclei were imaged in *z*-stack using a using a 405 nm laser to LSM980 confocal microscope (Zeiss). For scoring intestinal karyokinesis defects, images were converted into maximum intensity projections (MIPs) and scored. The quality was categorized as normal, or with defects based on its morphology. A normal score was given to distinct intestinal nuclei without any chromatin bridges or fusion; a defect indicated intestinal chromatin bridges or intestinal nuclei fusion.

### HUS-1::GFP foci formation

HUS-1::GFP transgenic worms [*daf-2(e1370);hus-1(op241);hus-1p::hus-1::gfp* and *daf-16(mgdf50);daf-2(e1370);hus-1(op241);hus-1p::hus-1::gfp*] were grown on control RNAi L1 onwards. At the late-L4 stage, the worms were subjected to IR (0 and 100 gy) and incubated for a duration of 6-8 or 24-27 hours at 20°C. The adults were subsequently mounted on microscope slides coated with a 2% agarose pad in 20 mM sodium azide, and z-stacked images were acquired using a 488 nm laser channel on a confocal microscope (LSM980 Zeiss). Under baseline conditions, HUS-1::GFP exhibits a diffuse presence in the cytoplasm and shows only weak localization in the chromatin of germ cells. However, in response to DNA damage (e.g., gamma radiation), HUS-1::GFP accumulates in nuclear foci across all germ cells, appearing as prominent bright foci. The bright HUS-1::GFP foci were quantified in 10 pachytene germ cells per gonad for each condition.

### Statistical tests

All statistical tests were performed utilizing the built-in functions of GraphPad Prism 10.1.0. In instances where we compared two conditions (continuous data), we applied a Student’s *t*-test with Welch’s correction, making no assumptions about consistent standard deviations. When comparing categorical data, we used the chi-square test. When comparing multiple conditions (continuous), we employed a Two-way ANOVA with Tukey’s multiple comparison test.

## Supporting information

C. elegans strains used and generated in the study

PCR primers used in the study

## Acknowledgements

We thank the members of the Molecular Ageing Laboratory, the Central Confocal Facility, and the Central Instrumentation Facility of the National Institute of Immunology. Some strains were provided by the CGC, which is funded by NIH Office of Research Infrastructure Programs (P40 OD010440).

## Funding

This project was partly funded by the National Bioscience Award for Career Development (BT/HRD/NBA/38/04/2016) and extramural grant (BT/PR27603/GET/119/267/2018) from the Department of Biotechnology, Government of India (https://dbtindia.gov.in/), Science and Engineering Research Board-Science and Technology Award for Research (SERB-STAR) award (STR/2019/000064), Jagadish Chandra Bose National Fellowship (JCB/2022/000021) and extramural grant (CRG/2022/000525) from the Ministry of Science and Technology, Government of India (https://serb.gov.in/page), as well as core funding from the National Institute of Immunology (to AM). GCS is supported by an ICMR SRF fellowship (RMBH/FW/2020/19), and UR by DBT-JRF fellowship DBT/2018/NII/1035. The funders had no role in study design, data collection and analysis, decision to publish, or preparation of the manuscript.

## Author Contributions

UR, GCS, AM conceptualized the project. UR, GCS, OD, RKR, RM and AG performed the experiments and analyzed data. UR, GCS, OD, AM wrote the manuscript. AM supervised the project and acquired funding.

## Competing Interest Statement

The authors disclose that there is no competing interest.

## Notes

### Competing Interest Statement

The authors have declared no competing interest.

