## Supplementary Figures for "DAF-16/FOXO maintains genome integrity following genotoxic stress"

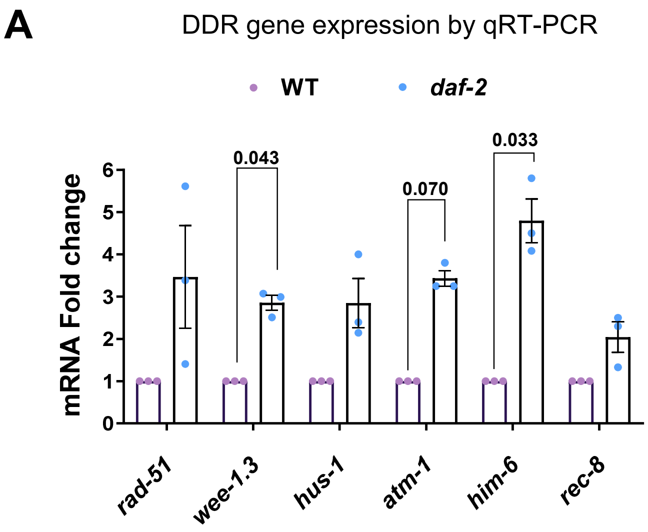


**Fig S1: DAF-16/FOXO regulates DNA damage repair pathway gene expression**

**(A)** Quantitative RT-PCR analysis showing mRNA fold change of DDR gene expression in wild-type (WT) and *daf-2(1370)* late-L4 staged worms grown on control RNAi. Expression levels were normalized to *actin*. The average of three biological replicates is shown. Unpaired *t*-test with Welch's correction.

Experiments were performed at 20°C. Source data are provided in Table 1.

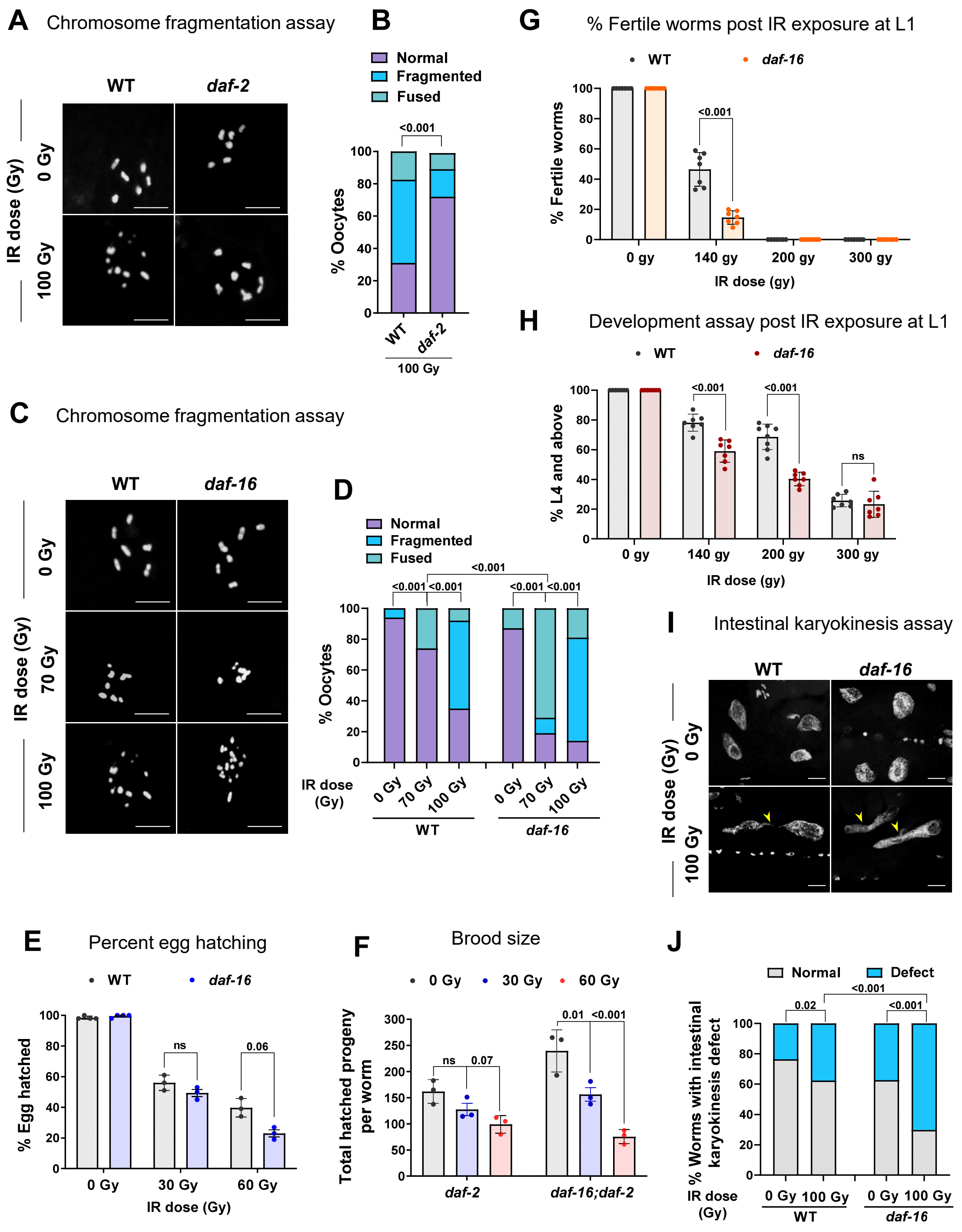
A
Chromosome fragmentation assay
WT
daf-2
IR dose (Gy)
0 Gy
100 Gy
B
Normal
Fragmented
Fused
<0.001
% Oocytes
100
80
60
40
20
0
WT
daf-2
100 Gy
G
% Fertile worms post IR exposure at L1
WT
daf-16
% Fertile worms
<0.001
0 gy
140 gy
200 gy
300 gy
IR dose (gy)
H
Development assay post IR exposure at L1
WT
daf-16
% L4 and above
<0.001
<0.001
ns
0 gy
140 gy
200 gy
300 gy
IR dose (gy)
C
Chromosome fragmentation assay
WT
daf-16
IR dose (Gy)
0 Gy
70 Gy
100 Gy
D
Normal
Fragmented
Fused
<0.001
<0.001 <0.001
<0.001 <0.001
% Oocytes
IR dose (Gy)
0 Gy
70 Gy
100 Gy
WT
daf-16
I
Intestinal karyokinesis assay
WT
daf-16
IR dose (Gy)
0 Gy
100 Gy
E
Percent egg hatching
WT
daf-16
% Egg hatched
ns
0.06
0 Gy
30 Gy
60 Gy
IR dose (Gy)
F
Brood size
0 Gy
30 Gy
60 Gy
Total hatched progeny per worm
300
250
200
150
100
50
0
ns
0.07
0.01
<0.001
daf-2
daf-16;daf-2
J
Normal
Defect
0.02
<0.001
<0.001
% Worms with intestinal karyokinesis defect
IR dose (Gy)
0 Gy
100 Gy
0 Gy
100 Gy
WT
daf-16

**Fig S2: DAF-16 maintains DDR efficiency in both germline and soma following genotoxic stress**

**(A)** Representative fluorescence images of DAPI-stained gonads (48 HPI) of WT and *daf-2(e1370)* worms grown on control RNAi, irradiated (0 and 100 Gy) at the late-L4 stage. Scale bars: 5 μm.

**(B)** Quantification for the degree of chromosome fragmentation as seen above. The oocyte chromosomes were categorized as normal or fragmented based on the number of oocyte chromosomes (normal = 6 chromosomes; fragmented = more than 6 chromosomes; fused = less than 6 chromosomes and fusion between two or more chromosomes). The combined data of three biological replicates (*n*≥64 oocytes for each replicate) is shown. Statistical comparisons between groups were performed using Chi-square analysis.

**(C)** Representative fluorescence images of DAPI-stained gonads (48 HPI) of WT and *daf-16(mgdf50)* worms grown on control RNAi, irradiated (0, 70, and 100 Gy) at the late-L4 stage. Scale bars: 5 μm**.**

**(D)** The quantification of the degree of chromosome fragmentation as seen above. The oocyte chromosomes were categorized as normal, fragmented, or fused based on morphology (normal = 6 chromosomes; fragmented = more than 6 chromosomes; fused = less than 6 chromosomes and fusion between two or more chromosomes). The average of three biological replicates (*n* ≥ 29) is shown. Statistical comparisons between groups were performed using Chi-square analysis.

**(E)** The percentage of hatched eggs of WT and *daf-16(mgdf50)* worms grown on control RNAi, irradiated (0, 30, and 60 Gy) at the late-L4 stage. Egg hatching was monitored for 12-24 HPI. The average of three biological replicates (*n* ≥ 100 per replicate) is shown. Unpaired *t*-test with Welch's correction

**(F)** The total number of hatched progenies in *daf-2(e1370)* and *daf-16(mgdf50);daf-2(e1370)* grown on control RNAi, irradiated (0, 30, and 60 Gy) at late-L4 stage. Average of three biological repeats (*n* ≥ 12 per replicate). Two-way ANOVA-Tukey's multiple comparisons test.

**(G)** The percentage of fertile worms in WT and *daf-16(mgdf50)* worms grown on control RNAi irradiated (0, 140, 200, and 300 Gy) at the L1 stage. The percentage of fertile worms (with eggs in the uterus) was subsequently quantified post-IR (when unirradiated worms reached the late day-1 stage). The average of seven biological replicates (*n*≥100 for each experiment) is shown. Unpaired *t*-test with Welch's correction.

**(H)** The percentage of larval development of WT and *daf-16(mgdf50)* worms grown on control RNAi irradiated (0, 140, 200, and 300 Gy) at the late-L4 stage. The percentage progressing to the L4 stage and beyond was subsequently quantified post-IR (when unirradiated worms reached the early day-1 stage). The average of seven biological replicates ($n \geq 100$ per replicate). Unpaired *t*-test with Welch's correction.

**(I)** Representative fluorescence images of DAPI-stained intestinal nuclei (48 HPI) of WT and *daf-16(mgdf50)* worms grown on control RNAi, irradiated (0 Gy and 100 Gy) at the late-L4 stage. The arrow points towards fused intestinal cell nuclei (intestinal karyokinesis defect). Scale bars: 10 μm.

**(J)** Quantification for the degree of intestinal karyokinesis defect as seen above. The quality was categorized as normal or with defects based on its morphology, where normal indicates distinct intestinal nuclei without any chromatin bridges or intestinal nuclei fusion, and defect indicates intestinal chromatin bridge or fused intestinal nuclei. The combined data of four biological replicates ($n \geq 124$ intestinal cells) is shown. Statistical comparisons between groups were performed using Chi-square analysis.

Experiments were performed at 20°C. Source data are provided in Table 1.

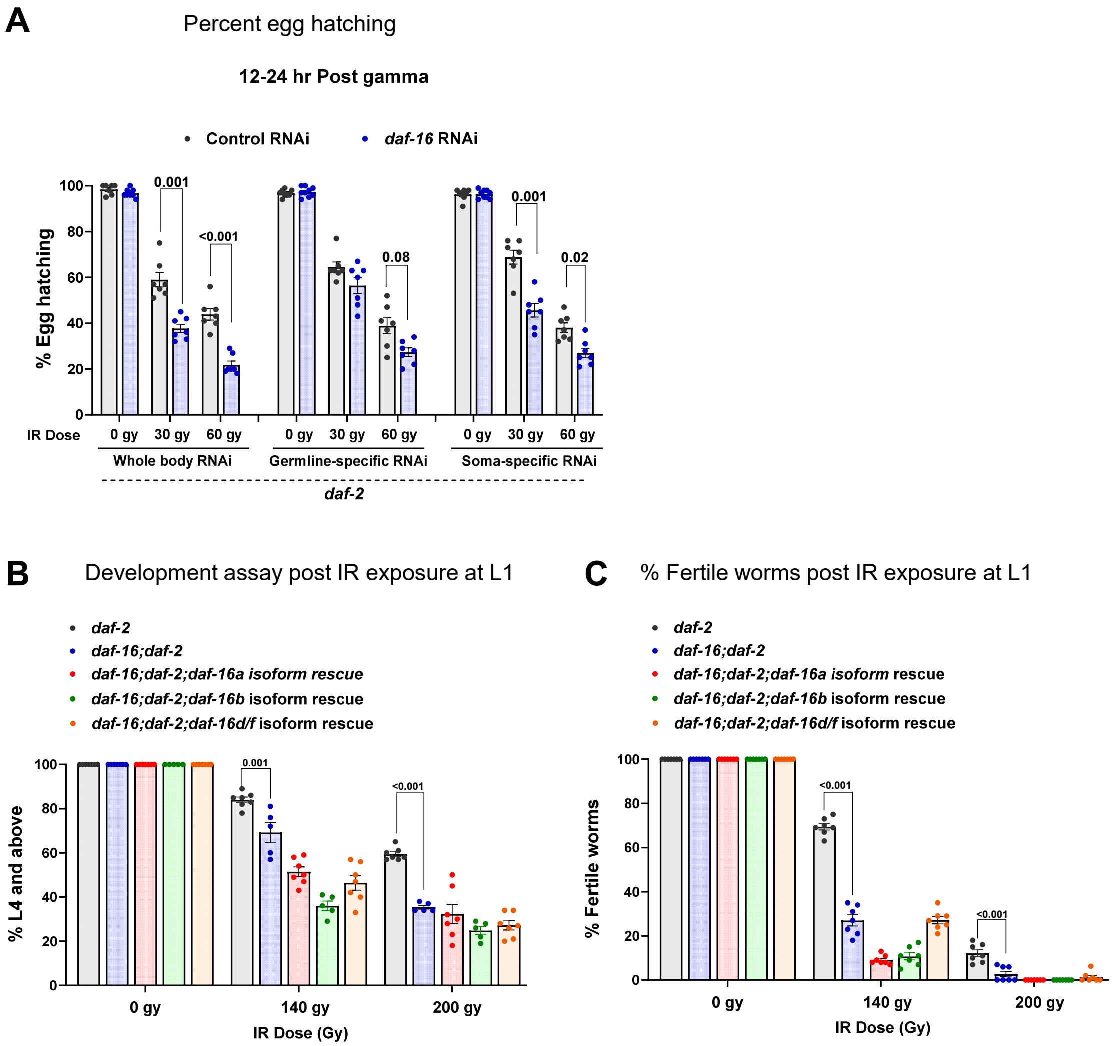

A
Percent egg hatching
12-24 hr Post gamma
Control RNAi
daf-16 RNAi
% Egg hatching
100
80
60
40
20
0
0.001
<0.001
0.08
0.001
0.02
IR Dose
0 gy
30 gy
60 gy
Whole body RNAi
Germline-specific RNAi
Soma-specific RNAi
daf-2
B
Development assay post IR exposure at L1
daf-2
daf-16;daf-2
daf-16;daf-2;daf-16a isoform rescue
daf-16;daf-2;daf-16b isoform rescue
daf-16;daf-2;daf-16d/f isoform rescue
% L4 and above
0.001
<0.001
0 gy
140 gy
200 gy
IR Dose (Gy)
C
% Fertile worms post IR exposure at L1
daf-2
daf-16;daf-2
daf-16;daf-2;daf-16a isoform rescue
daf-16;daf-2;daf-16b isoform rescue
daf-16;daf-2;daf-16d/f isoform rescue
% Fertile worms
<0.001
<0.001
0 gy
140 gy
200 gy
IR Dose (Gy)

**Fig S3: DAF-16 is required in both germline and soma to maintain germline DDR efficiency.**

**(A)** The percentage of hatched eggs of *daf-2(e1370)*, *daf-2*;*rde-1(mkc36);sun-1p::rde-1* (germline-specific RNAi), and *daf-2*;*ppw-1(pk1425)* (soma-specific RNAi) worms grown on control or *daf-16* RNAi, irradiated (0, 30, and 60 Gy) at the late-L4 stage. Egg hatching was monitored for 12-24 HPI. The average of seven biological replicates ($n \geq 12$ per replicate). Unpaired *t*-test with Welch's correction.

**(B)** The percentage of larval development of *daf-2(e1370), daf-16(mgdf50);daf-2(e1370)*, and different DAF-16 isoforms rescued *(daf-16a::RFP, daf-16b::CFP, and daf-16d/f::GFP)* in *daf-16(mgdf50);daf-2(e1370)* worms grown on control RNAi, irradiated (0, 140, and 200 Gy) at the late-L4 stage. The percentage progressing to the L4 stage and beyond was subsequently quantified post-IR (when unirradiated worms reach the early day-1 stage). The average of seven biological replicates ($n \geq 100$ per replicate) is shown. Two-way ANOVA with Tukey's multiple comparison test.

**(C)** The percentage of fertile worms in *daf-2(e1370), daf-16(mgdf50);daf-2(e1370)* and different DAF-16 isoforms rescued *(daf-16a::RFP, daf-16b::CFP, and daf-16d/f::GFP)* in *daf-16(mgdf50);daf-2(e1370)* worms grown on control RNAi, irradiated (0, 140, and 200 Gy) at the late-L4 stage. The percentage of fertile worms (with eggs in the uterus) was subsequently quantified post-IR (when unirradiated worms reached the late day-1 stage). The average of seven biological replicates ($n \geq 100$ for each experiment). Two-way ANOVA with Tukey's multiple comparison test.

Experiments were performed at 20°C. Source data are provided in Table 1.

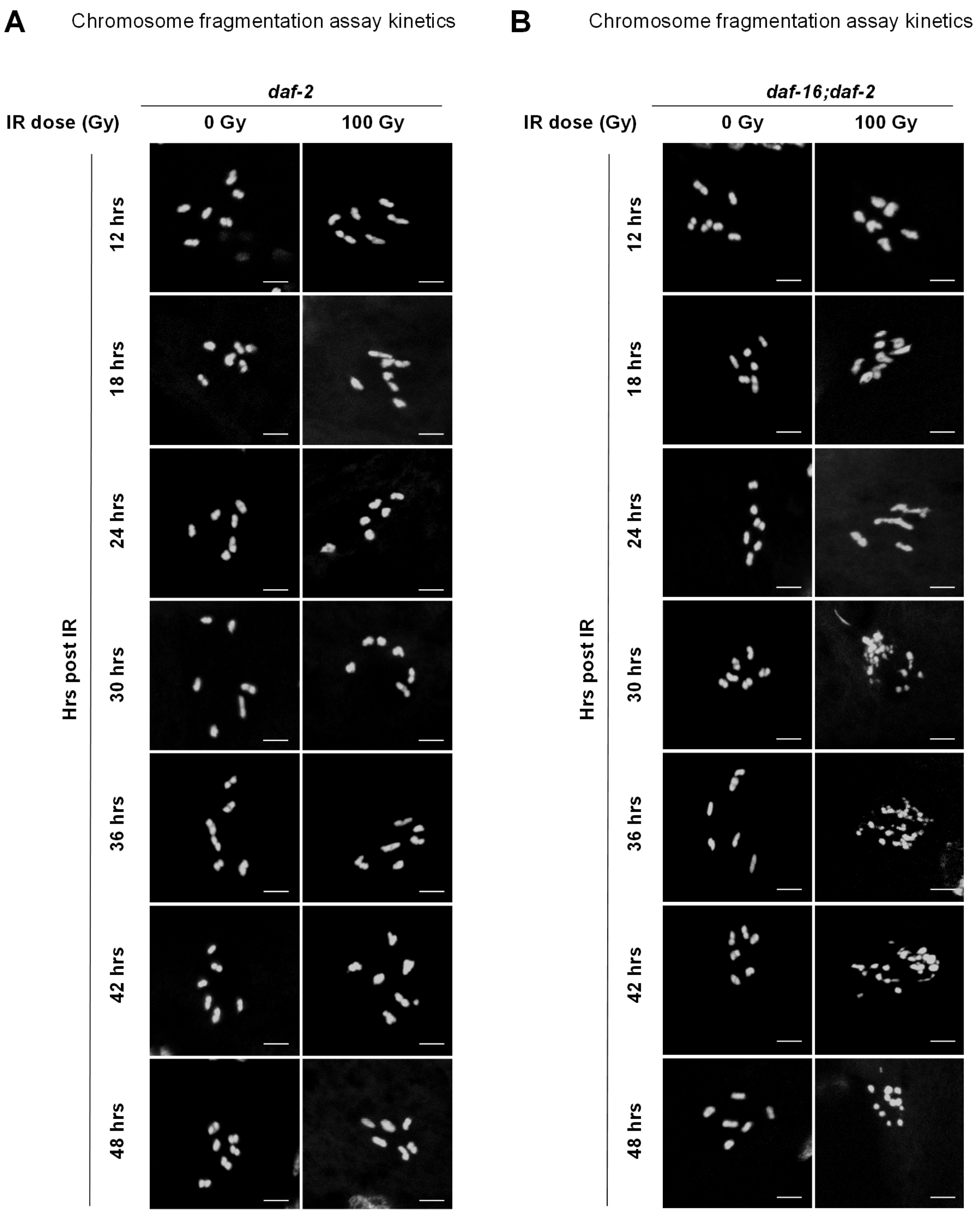

A
Chromosome fragmentation assay kinetics
daf-2
IR dose (Gy)
0 Gy
100 Gy
Hrs post IR
12 hrs
18 hrs
24 hrs
30 hrs
36 hrs
42 hrs
48 hrs
B
Chromosome fragmentation assay kinetics
daf-16;daf-2
IR dose (Gy)
0 Gy
100 Gy
Hrs post IR
12 hrs
18 hrs
24 hrs
30 hrs
36 hrs
42 hrs
48 hrs

**Fig S4: Kinetics of DNA damage repair is accelerated upon DAF-16 activation**.

**(A, B)** Representative fluorescence images of DAPI-stained gonads (12,18, 24, 30, 36, 42, and 48 HPI) of *daf-2(e1370)* and *daf-16(mgdf50)*;*daf-2(e1370)* worms grown on control RNAi, irradiated (0 and 100 Gy) at the late-L4 stage. Scale bars 5 µm.

Experiments were performed at 20°C.

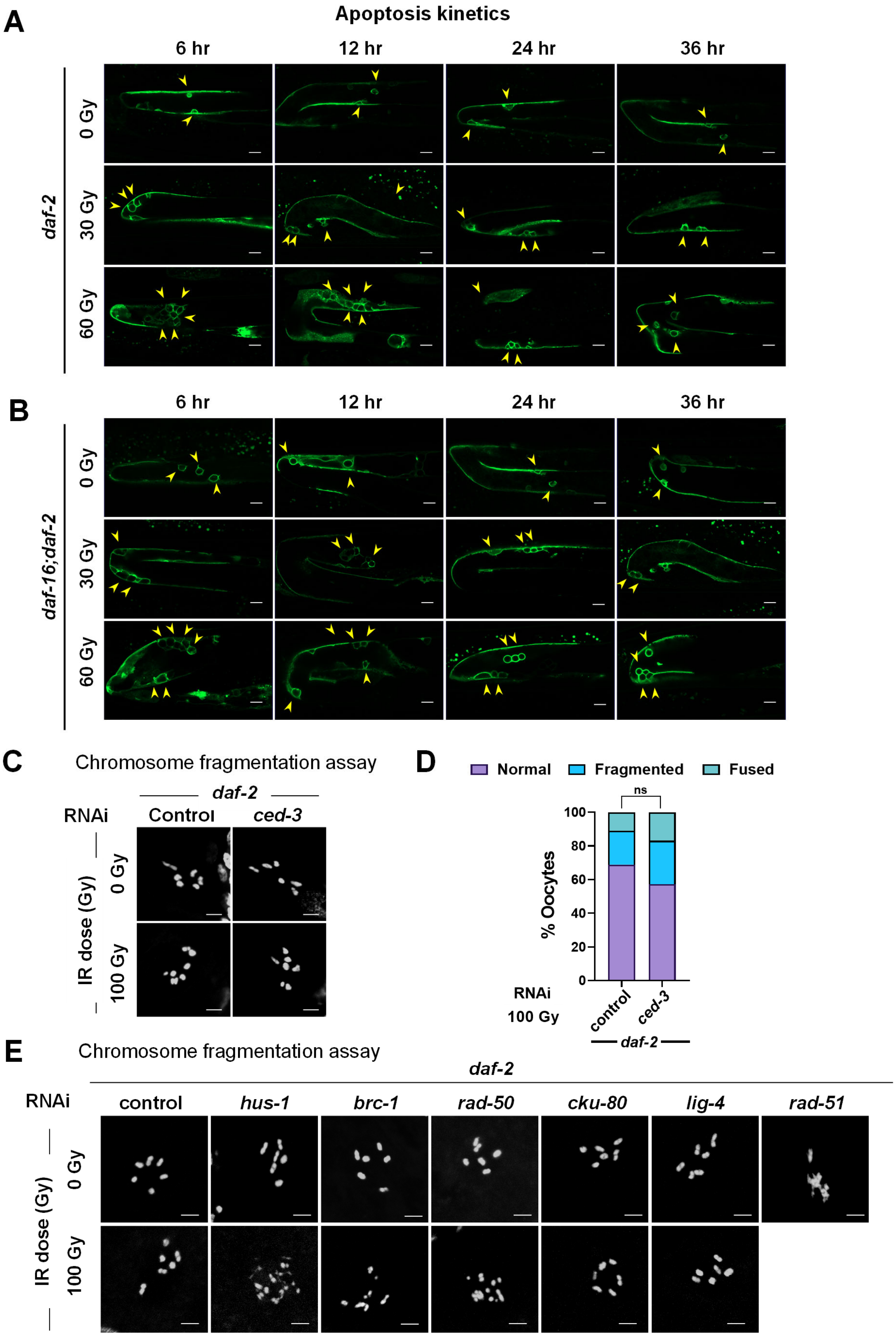
A
Apoptosis kinetics
6 hr
12 hr
24 hr
36 hr
daf-2
0 Gy
30 Gy
60 Gy
B
daf-16;daf-2
C
Chromosome fragmentation assay
daf-2
RNAi
Control
ced-3
IR dose (Gy)
100 Gy
D
Normal
Fragmented
Fused
ns
% Oocytes
100
80
60
40
20
0
RNAi
control
100 Gy
E
hus-1
brc-1
rad-50
cku-80
lig-4
rad-51

**Fig S5: Activated DAF-16 is dependent on the canonical DDR components, but not apoptosis, to enhance germline DDR efficiency**

**(A, B)** Representative images showing the kinetics of apoptosis (6, 12, 24, and 36 HPI) in the gonadal arm of *daf-2(e1370);ced-1::GFP* (A) and *daf-16(mgdf50);daf-2(e1370);ced-1::GFP* (B) worms grown on control RNAi, irradiated (0, 30, and 60 Gy) at the late-L4 stage**.** Scale bars: 10 μm.

**(C)** Representative fluorescence images of DAPI-stained gonads (48 HPI) of *daf-2(e1370)* worms grown for two generations on control and *ced-3* RNAi, irradiated (0 and 100 Gy) at the late-L4 stage. Scale bars: 5 μm.

**(D)** The quantification of the degree of chromosome fragmentation as seen above. The oocyte chromosomes were categorized as normal, fragmented, or fused based on their morphology (normal = 6 chromosomes; fragmented = more than 6 chromosomes; fused = less than 6 chromosomes and fusion between two or more chromosomes). Combined data from three biological replicates ($n \geq 36$) are presented. Statistical comparisons between groups were performed using Chi-square analysis.

**(E)** Representative fluorescence images of DAPI-stained gonads (48 HPI) of *daf-2(e1370)* worms grown for two generations on control, *hus-1, rad-50*, *cku-80*, *lig-4*, and *rad-51* RNAi (two-generational RNAi), irradiated (0 and 100 Gy) at the late-L4 stage. Scale bars: 5 μm.

Experiments were performed at 20°C. Source data are provided in Table 1.
